# Bacterial “galactosemia” is caused by cytoplasmic interference of an essential cell wall biosynthesis glycosyltransferase

**DOI:** 10.1101/2021.01.22.426804

**Authors:** Cameron Habib, Anna Mueller, Ming Liu, Jun Zhu, Tanja Schneider, Ethan Garner, Yingjie Sun, Yunrong Chai

## Abstract

Bacterial galactosemia or “galactose death,” triggered by incomplete galactose metabolism, was first discovered in *Escherichia coli* and *Salmonella* six decades ago, and later in many other microorganisms, yet the mechanism for the toxicity and subsequent cell death remains unclear. In *Bacillus subtilis,* galactosemia is manifested by a buildup of uridine-diphosphate-galactose (UDP-Gal) and a strong toxicity phenotype characterized by cell shape abnormality and rapid cell lysis. Here we present evidence that in *B. subtilis,* the toxicity is due to inhibition of cell wall biosynthesis through interference of the essential glycosyltransferase MurG that carries out lipid II synthesis from lipid I and uridine-diphosphate-*N*-acetyl-glucosamine (UDP-GlcNAc). Single-molecule imaging reveals real-time inhibition of cell wall biosynthesis and MurG activities in cells exhibiting toxicity. We further show that *in vitro*, MurG is able to utilize UDP-Gal as a substrate generating a “toxic” lipid II, causing a potential poisoning effect on peptidoglycan crosslinking. Evidence also suggests a similar mechanism in *Vibrio cholerae* and *Staphylococcus aureus*. Lastly, a strong synergistic lethality was seen in *S. aureus* wild-type cells treated with both galactose and sub-lethal doses of cell-wall antibiotics. Our study provides mechanistic explanation of the toxicity associated with bacterial galactosemia and its potential application in antibacterial solutions.

**Significance:** Galactosemia is a potentially fatal genetic disorder due to incomplete galactose metabolism, found in both eukaryotic and prokaryotic organisms. The molecular mechanisms of galactosemia-associated toxicity remain unclear in all cases. Here we present evidence that in the bacterium *Bacillus subtilis*, the toxicity is due to interference of an essential glycosyltransferase, MurG, which concerts lipid I to lipid II during peptidoglycan biosynthesis, by a nucleotide sugar derived from galactose metabolism. This interference leads to a halt of cell wall biosynthesis and structural defects causing rapid cell lysis. Our evidence also suggests a similar mechanism in other bacteria such as *Staphylococcus aureus* and *Vibrio cholerae.* Our study may help solve the long-time puzzle of bacterial galactosemia first uncovered six decades ago.

## Introduction

Galactosemia arises from mutation or deficiency in the near ubiquitous Leloir pathway ^1^. This pathway, conserved in both prokaryotes and eukaryotes, consists of three enzymes (GalK, GalT, and GalE) and is responsible for the metabolism of α-D-galactose to glucose-1-phosphate (Fig. 1A). The metabolic process begins as galactokinase (GalK) converts α-D-galactose to galactose-1-phosphate (Gal-1-P), followed by galactose-1-P-uridylyltransferase (GalT) converting Gal-1-P and uridine-diphosphate-glucose (UDP-Glc) to uridine-diphosphate-galactose (UDP-Gal) and glucose-1-P. Lastly, UDP-Gal is converted back to UDP-Glc by UDP-galactose 4-epimerase (GalE) (Fig. 1A)^2^. The metabolism from α-D-galactose to glucose-1-P is believed to be self-sustainable by the activities of these three enzymes ^1^. In bacteria, the inability to completely metabolize galactose causes a strong toxicity phenotype, a phenomenon phrased as “galactose death” or “bacterial galactosemia,” first described in *E. coli* and *Salmonella* in 1959 ^3,4^ and later observed in many other bacteria ^5,6^, with the underlying mechanism still unknown.

**Figure 1.**
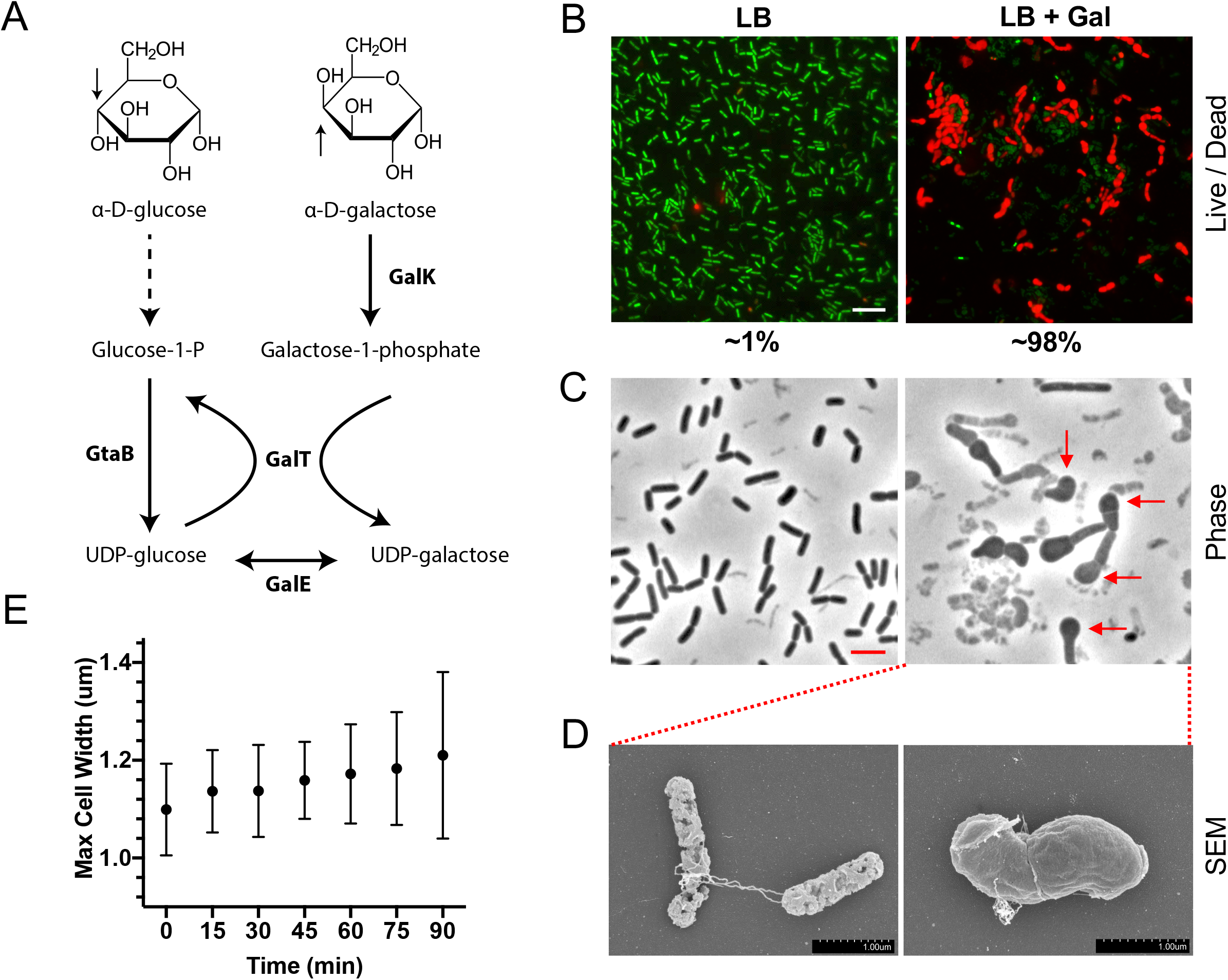
Incomplete galactose metabolism causes cell shape deformation and cell lysis in *B. subtilis*. **(A)** An overview of the Leloir pathway in *B. subtilis*. Galactose metabolism is carried out by three enzymes, galactokinase (GalK), which first catalyzes α-D-galactose to galactose-1-phosphate, followed by galactose-1-phosphate uridylyl-transferase (GalT), which transfers a uridine-diphosphate group from UDP-glucose to galactose-1-phosphate, forming UDP-galactose and glucose-1-phosphate. Finally, UDP glucose 4-epimerase (GalE) catalyzes the reversible reaction from UDP-galactose to UDP-glucose. In addition, a UTP-glucose-1-phosphate uridylyl-transferase (GtaB) replenishes the pool of UDP-glucose by catalyzing the reaction from glucose-1-phosphate to UDP-glucose. The two sugar epimers, α-D-Glucose and α-D-galactose, differ only in the orientation of the hydroxyl group at C4 carbon (indicated by arrows). **(B)** The Δ*galE* mutant of *B. subtilis* (CH62) was grown in LB and LB+galactose (0.5%, w/v) to log phase while shaking at 37°C. Cells were collected, stained for live (green) / dead (red) fluorescence dyes, and imaged. The percentage of live / dead cells was quantified (shown below panels). Scale bar is 10 μm. **(C)** Cells were imaged under phase contrast microscopy to determine the effect of the toxicity on cell morphology. Structural defects (red arrows) including bulges, rounding out of bacilli shape, and cell lysis were observed. Scale bar is 4 μm. **(D)** Scanning electron microscopy shows structural defects of cells, which ranged from shape deformities to complete loss of integrity. Scale bar is 1 μm. **(E)** After addition of galactose the progression of toxicity was measured over time as a function of maximal cell width of ~100 cells using microscopy. Increasing maximal width over time indicates more severe structural defects associated with incomplete galactose metabolism. Error bars represent standard deviations.

In *Bacillus subtilis*, a deletion in the *galE* gene causes a buildup of UDP-Gal when cells grow in excess galactose and a strong toxicity phenotype characterized by cell shape deformation and rapid cell lysis ^6,7^. While buildup of UDP-Gal causes toxicity, this nucleotide sugar is needed for a number of important processes, including synthesis of an exopolysaccharide (EPS) for biofilm formation in *B. subtilis* ^7,8^. Therefore, producing UDP-Gal but maintaining its cellular level below a threshold that triggers toxicity must be an important regulation during galactose metabolism. In a previous work, we uncovered a shunt mechanism in *B. subtilis* allowing bypass of the toxicity upon strong biofilm induction when UDP-Gal was used as a precursor for EPS biosynthesis and exported in large quantity ^7^. Galactose can be naturally and abundantly available to *B. subtilis* when this soil bacterium utilizes complex carbohydrates available in the environment such as plant-derived galactan ^9^. *B. subtilis* possesses a dedicated pathway for uptake and hydrolysis of galactan into galactose for metabolic purposes as well as to make EPS for biofilm formation. Biofilms in turn play a key role in establishing intimate relationships between *B. subtilis* and the plant ^7,8,10–13^.

Bacterial cell wall consists of cross-linked peptidoglycan chains, whose building block is delivered via a membrane lipid carrier, lipid II. Synthesis of lipid II is carried out in the cytoplasm by an essential glycosyltransferase MurG, through the addition of *N*-acetyl-glucosamine (GlcNAc) to lipid I, by utilizing a nucleotide sugar UDP-*N*-acetyl-glucosamine (UDP-GlcNAc) ^14,15^. Once synthesized, lipid II is flipped across the cytoplasmic membrane by a flippase, and the sugar-peptide building block polymerized into full-length peptidoglycan by extracellular penicillin-binding proteins ^16–18^. The biosynthetic machinery responsible for peptidoglycan polymerization in both *E. coli* and *B. subtilis* has been shown to be associated with MreB filaments, as they move around the cell, a circumferential movement driven by, and directly correlated to, peptidoglycan synthesis ^19,20^. Using fluorescent protein fusions to MreB, the movement of MreB filaments can be visualized under Total Internal Reflection Fluorescence (TIRF) microscopy and has been used as an assay for measuring dynamic cell wall biosynthesis ^21^.

In this study, we investigated the mechanism of cell toxicity associated with bacterial galactosemia in *B. subtilis*. Our evidence suggests a strong link between the toxicity and inhibition of cell wall biosynthesis, with the essential glycosyltransferase MurG and lipid precursor biosynthesis as targets.

## Results

### Galactose triggers cell shape abnormality and rapid cell lysis in a *B. subtilis* Δ*galE* mutant

*B. subtilis* Δ*galE* cells quickly lyse when grown in the presence of galactose ^7^. Evidence suggests that this toxicity is due to accumulation of UDP-Gal as a result of incomplete galactose metabolism (Fig. 1A)^6^. To further characterize the toxicity, Δ*galE* cells were grown in LB supplemented with galactose (0.5%, w/v), periodically collected, stained with live/dead fluorescent dyes, and imaged under fluorescent microscopy. We found that cell optical density (OD_600_) of the culture peaked at around 0.3, and an hour past the peak, the ratio of dead versus total cells increased exponentially to approximately 98% based on live/dead cell staining (Fig. 1B). In the absence of galactose, this number was approximately 1%. Phase contrast imaging showed gross structural deformities of cells such as rounding out of the regular bacilli shape, cell wall bulges, and cell lysis not seen in untreated cells (indicated by arrows, Fig. 1C). Scanning electronic microscopy showed clear damage in cell structural integrity, with deformations of cell shape and apparent defects in the cell envelope (Fig. 1D). To quantitatively analyze how individual cells were impacted, the maximum cell width was determined by single cell analysis under microscopy and the distribution plotted. 90 min after addition of galactose, the average width distribution of the galactose-treated cells increased ~20% over that of untreated cells (Fig. 1E), indicating that both the ratio and the severity of cells exhibiting toxicity increased over time. In the same assays without addition of galactose, the doubling time of Δ*galE* cells did not differ from that of the WT (Fig. S1A).

### Production, not depletion, of UDP-glucose, contributes to toxicity

Uridine-diphosphate-glucose (UDP-Glc) is one of the intermediates during galactose metabolism (Fig. 1A). It is also a common precursor for syntheses of complex polysaccharides such as teichoic acids, lipopolysaccharides, and a signal for cell size control in bacteria ^22,23^. Given the importance of UDP-Glc, it was previously proposed that the toxicity associated with incomplete galactose metabolism was due to depletion of UDP-Glc, rather than accumulation of UDP-Gal ^24^. Depletion of UDP-Glc can occur when the uridine-di-phosphate group is transferred from UDP-Glc to Gal-1-P to form UDP-Gal and Glu-1-P (Fig. 1A). In cells lacking GalE, regeneration of UDP-Glc from UDP-Gal is blocked, thus gradually depleting UDP-Glc while accumulating UDP-Gal ^6^. In the absence of active galactose metabolism, the primary source of UDP-Glc is believed to be made from Glc-1-P by a UDP-glucose pyrophosphorylase (GtaB, Fig. 1A)^22,25^. In *B. subtilis*, loss of GtaB activity leads to depletion of UDP-Glc ^25^. To assess the impact of UDP-Glc depletion in *B. subtilis*, a *gtaB* deletion strain was created and cells were grown in LB without addition of galactose. No clear growth defect or toxicity was observed (Fig. 2A). Although not essential for growth, the Δ*gtaB* mutant was found completely deficient in biofilm formation (Fig. S1B). We previously showed that the EPS of the *B. subtilis* biofilm matrix was rich in glucose ^7^. The biofilm deficiency of the Δ*gtaB* mutant suggests that UDP-Glc is the precursor for providing the glucose moiety of the EPS. It also confirms that GtaB activity is the primary source of UDP-Glc synthesis in *B. subtilis*.

**Figure 2.**
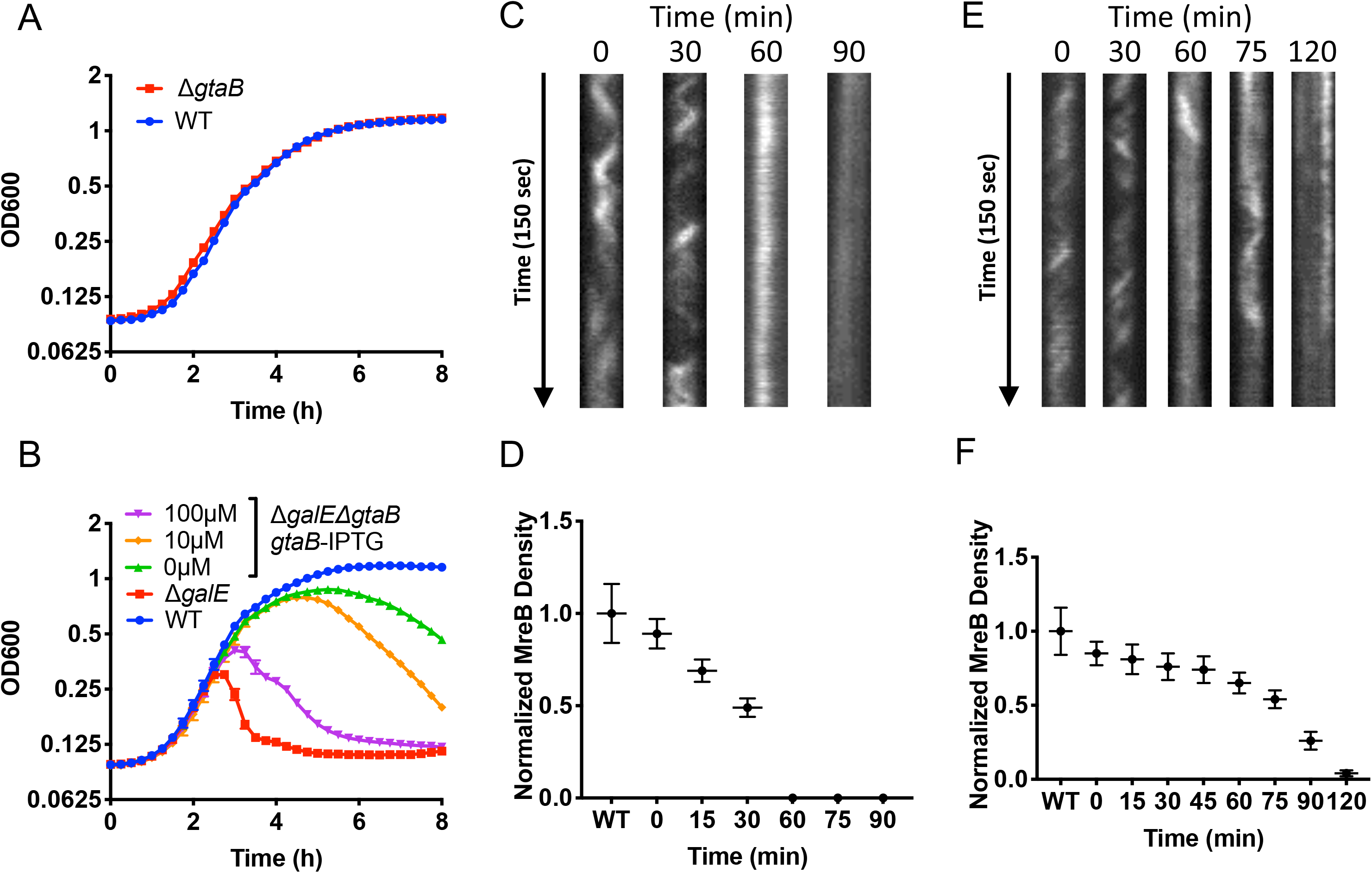
(A-B) GtaB is required for toxicity. **(A)** WT *B. subtilis* (3610) and a strain deficient in *gtaB* (YC876) were grown in LB shaking at 37°C and OD_600_ values of the cultures were measured every 15 minutes for a period of 8 hours. Representative biological replicate shown with error bars representing standard deviation. **(B)** An IPTG inducible copy of *gtaB* was introduced into the Δ*galE*Δ*gtaB* double mutant and the result strain (CH117) was grown in shaking LB with galactose (0.5%, w/v) and varying amounts of IPTG for *gtaB* induction, together with the WT (3610) and the Δ*galE* mutant (CH62). As judged by the OD_600_ values of the cultures. Representative biological replicate shown with error bars representing standard deviation. **(C-F) Toxicity is due to inhibition of cell wall biosynthesis. (C-D)** Kymograph of MreB movement in the Δ*galE* mutant. The polymerization and movement of MreB filaments is tracked using MreB-mNeonGreen fusions, a motion dependent upon active cell wall synthesis. Δ*galE* cells expressing the MreB-mNeonGreen fusion protein were immobilized on a CH agar pad with galactose (0.5%, w/v) and imaged with time lapse TIRF every 30 minutes. The cellular MreB-mNeonGreen location was recorded **(C)**, analyzed, then plotted **(D)**. The number of MreB-mNeonGreen filaments was found to substantially decrease over time, dissolving completely 60 minutes after galactose exposure. **(E-F)** Kymograph of MreB movement in the Δ*galE* mutant with *murG* overexpression (1 mM IPTG). **(E)** MreB filament formation and movement was recorded similarly in the Δ*galE* cells overexpressing *murG* by growth on agar pad containing galactose (0.5%, w/v) and 500 μM IPTG. **(F)** The number of MreB filaments was found partially restored by *murG* overexpression in 60, 75, and 90 min samples.

Upon further investigation, we found that not only does UDP-Glc depletion have no impact on cell viability, but production of UDP-Glc by GtaB is a prerequisite for accumulating UDP-Gal and causing toxicity during galactose metabolism. A double mutant of Δ*galE*Δ*gtaB* with an IPTG-inducible copy of *gtaB* was constructed. The engineered strain was grown in LB with galactose and varying amounts of IPTG to induce *gtaB* expression. The observed toxicity, judged by growth curve, correlated well with *gtaB* induction in a dose dependent manner (Fig. 2B). These findings suggest that continuous production of UDP-Glc by GtaB is necessary for accumulating UDP-Gal and triggering toxicity in the Δ*galE* cells (Fig. 1A).

### Toxicity is associated with inhibition in cell wall biosynthesis

Given the cell shape abnormality observed in the Δ*galE* cells and their resemblance to known cell shape phenotypes caused by defects in cell wall biosynthesis ^26^, we sought to test if the toxicity is linked to impaired cell wall biosynthesis. Cell wall biosynthetic machinery has been shown to be bound to and pull along MreB filaments as cargo ^19,21,27^, a motion driven by cell wall synthesis. Inhibition of cell wall synthesis via cell wall targeting antibiotics can causes MreB movement to cease ^19,28^. To determine in real-time if the dynamics of cell wall biosynthesis were altered in cells exhibiting toxicity, a MreB-mNeonGreen fusion was expressed in the *B. subtilis* Δ*galE* mutant. Cells were immobilized under a CH agar pad with galactose (0.5%, w/v) and the activity of the MreB filaments observed using Total Internal Reflection Fluorescence (TIRF) microscopy with imaging every 30 minutes. Movement of the fluorescent foci in single cells over the course of the time-lapse imaging was calculated and plotted (Fig. 2C, D, and Supple. Movie 1). The results indicate that movement of the MreB filaments decreased dramatically over time, with movement ceasing and filaments depolymerizing completely ~60 minutes after galactose addition (Fig. 2C, D). These results suggest that cell wall biosynthesis was blocked in the Δ*galE* cells exhibiting toxicity.

### Toxicity is not likely due to inhibition in biosynthesis of UDP-GlcNAc

Cell wall biosynthesis initiates in the cytoplasm by diverting the glycolytic sugar fructose-6-phosphate (Fru-6-P) to glucosamine-6-phosphate (GlcN-6-P) by a glutamine-fructose-6-phosphate aminotransferase (GlmS, Fig. 3A). A recent study showed a similar cell shape deformation and lysis phenotype, as seen in Δ*galE*, in a *B. subtilis* Δ*glmR* mutant ^29^. *glmR* encodes a regulatory protein for the aminotransferase GlmS, which carries out the first step in the synthesis of UDP-GlcNAc (Fig. 3A). When the cellular UDP-GlcNAc levels are low, GlmR activates GlmS, thus promoting synthesis of UDP-GlcNAc. In the presence of excess UDP-GlcNAc, GlmR is inhibited upon binding directly to UDP-GlcNAc. This serves as a checkpoint mechanism for UDP-GlcNAc biosynthesis (Fig. 3A)^29^.

**Figure 3.**
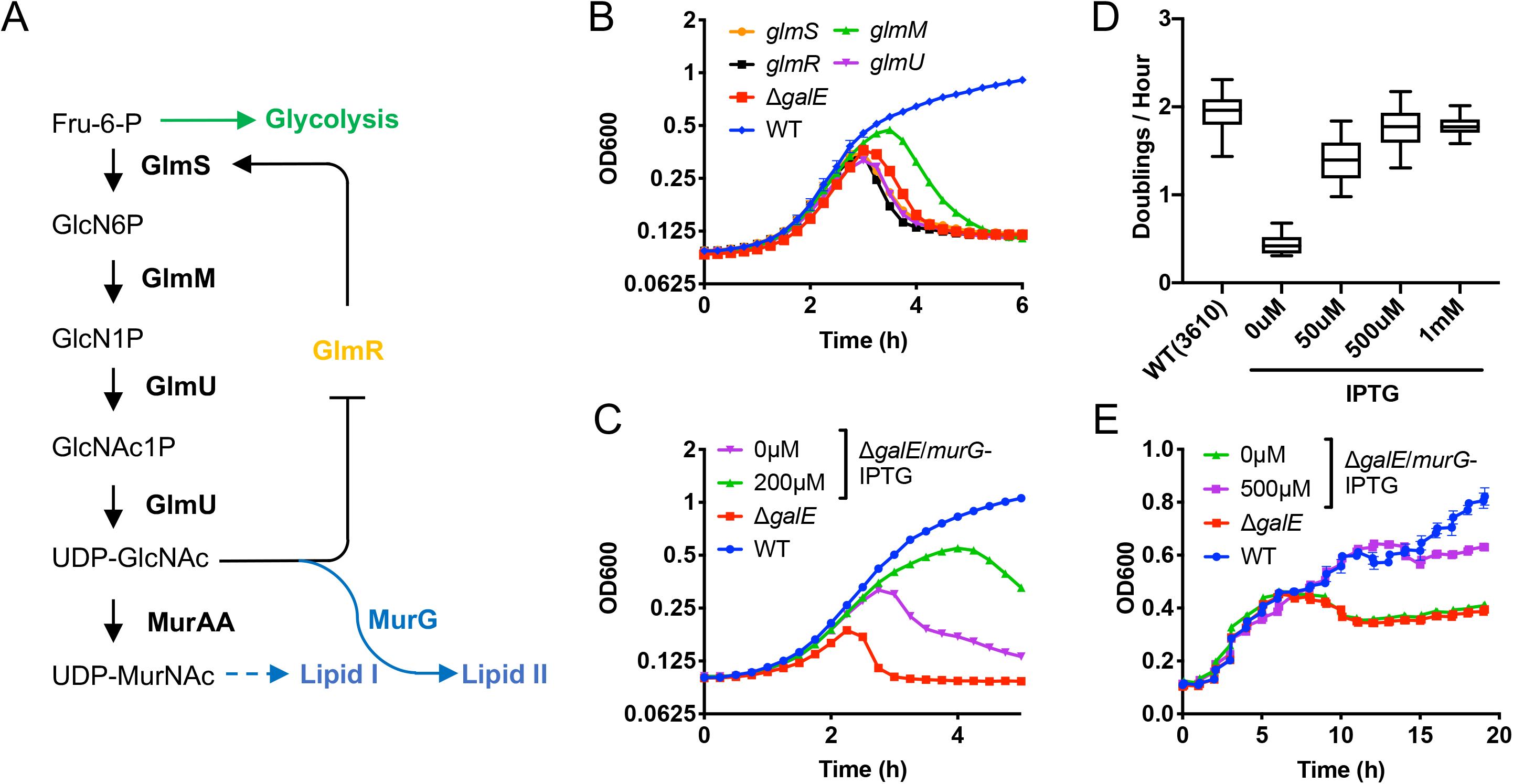
(A-B) Toxicity is not likely due to inhibition in synthesis of UDP-GlcNAc. **(A)** The biosynthetic pathway of UDP-GlcNAc and UDP-MurNAc. GlmS, GlmM, GlmU, MurAA are enzymes that carry out specific reactions from Fru-6-P to UDP-GlcNAc and UDP-MurNAc. GlmR is a regulatory protein for GlmS and the activity of GlmR is negatively regulated by UDP-GlcNAc. **(B)** The Δ*galE* mutant bearing either an IPTG-inducible copy of *glmS*, *glmR*, *glmM*, or *glmU* involved in UDP-GlcNAc biosynthesis was grown in LB shaking broth in a plate reader. OD_600_ of the cultures was measured periodically. Representative biological replicate shown with error bars representing standard deviation. **(C-E) MurG is a possible direct target of inhibition in cells exhibiting toxicity. (C)** An IPTG inducible copy of *murG* was introduced into the *B. subtilis* Δ*galE* mutant (CH112) and grown in LB shaking at 37°C with or without galactose (0.5%, w/v) and 0.2 mM IPTG, along with the WT and Δ*galE* strains in LB with galactose. OD_600_ values were measured every 15 minutes. Representative biological replicate shown with error bars representing standard deviation. **(D)** The effect of *murG* overexpression on growth rate and toxicity rescue at the single cell resolution. The Δ*galE murG* overexpression strain (CH112) was plated on a CH agar pad containing galactose (0.5%, w/v) and 0, 50, 500, and 1000 μM of IPTG and imaged every 15 minutes to determine growth rate as a function of doublings per hour. WT cells were used as a control. **(E)** A similar *murG* overexpression strain was constructed in the *V. cholerae* Δ*galE* mutant using its native *murG* gene. Cells were grown in LB shaking broth and OD_600_ values were measured periodically. *murGVC* was overexpressed upon addition of 500 μM IPTG. Strains used for the *V. cholerae* assay include WT(C6706), Δ*galE*(VCA0774), and Δ*galE*-*murG*^OV^(VC2401). Error bar represents standard deviation.

Since most enzymes involved in UDP-GlcNAc biosynthesis are essential, we hypothesized that if UDP-Gal targets any of these enzymes, it can cause toxicity. For example, it is possible that UDP-Gal directly binds to and inhibits GlmR and thus blocks UDP-GlcNAc biosynthesis given that GlmR also binds to UDP-Glc, which is structurally similar to UDP-Gal ^30^. If so, providing GlmR in excess could partially alleviate toxicity. We took a genetic approach to test this by overexpressing *glmR* in the Δ*galE* mutant. However, no toxicity rescue was observed (Fig. 3B). *glmS* was similarly overexpressed since it was shown that the cell shape deformation and lysis phenotype of Δ*glmR* can be rescued by overexpression of *glmS* ^29^. Again, no rescue in toxicity was observed (Fig. 3B). Finally, we overexpressed *glmU* and *glmM*, respectively, and none showed toxicity rescue (Fig. 3B). Thus, our genetic evidence suggests that biosynthesis of UDP-GlcNAc, the initial cytoplasmic stage in peptidoglycan biosynthesis, is likely not the target associated with toxicity.

### Overexpression of *murG* partially suppresses the toxicity

The final step in lipid II biosynthesis is carried out by the essential glycosyltransferase MurG, through the addition of *N*-acetyl-glucosamine to lipid I (Fig. 3A). To test if MurG is a possible target, we introduced an IPTG inducible copy of *murG* into the *B. subtili*s Δ*galE* mutant. The resulting strain was grown in the presence of both galactose and IPTG. Upon addition of 200 μM IPTG, there was a significant reduction of toxicity (as judged by the growth curve) compared to no IPTG addition (Fig. 3C). Note that a slight toxicity reduction was seen even in the absence of IPTG, likely due to leaky expression of *murG* from the *hyperspank* promoter. This suggests that *murG* overexpression can partially rescue the toxicity phenotype in *B. subtilis*. This rescue was unique to MurG, as other essential enzymes *mnaA, tagB*, and *murAA*, also involved in the utilization of UDP-GlcNAc, did not show any rescue upon similar overexpression (Fig. S1C).

We further assayed the toxicity rescue by *murG* overexpression by analyzing the growth of single cells. The Δ*galE* cells with the inducible *murG* were grown on a CH agar pad containing galactose (0.5%, w/v) and varying concentrations of IPTG and imaged to determine the doubling time of individual cells. Strong *murG* expression (addition of 500 μM IPTG) caused cells to regain close to the WT growth rates (Fig. 3D). Further, cells were found to maintain a consistent max cell width for a greater period of time as compared to without MurG overexpression (Fig. S1D). As the control, *murG* overexpression did not result in an increased growth rate in the absence of galactose (Fig. S1A, S1E). To determine if *murG* overexpression indeed relieved inhibition of cell wall biosynthesis, the *mreB-mNeonGreen* construct previously utilized was introduced into the above *murG* inducible strain and the resulting strain was similarly imaged for MreB-mNeonGreen movement and filamentation. Upon *murG* overexpression, the movement and number of MreB filaments increased significantly above those in Δ*galE* cells, at 60 and even 90 min after addition of galactose (Fig. 2E-F, compare Fig. 2D and 2F; Supple. Movie 2). Taken together, this suggests that MurG activity may be the target or cause of UDP-Gal-mediated toxicity. Lastly, overexpression of neither MurAA, the first committed step in peptidoglycan synthesis ^31^, nor MraY, the first membrane step of peptidoglycan synthesis in which lipid I is generated ^32^, resulted in rescue on blocked cell wall biosynthesis as observed upon overexpression of *murG* (Supple. Movie 3).

### *In vivo* evidence for inhibition of peptidoglycan precursor synthesis and MurG in cells exhibiting toxicity

Inhibition of MurG or subsequent cell wall biosynthesis step can cause an accumulation of lipid precursors, and ultimately halt of cell wall biosynthesis ^15,33,34^, which can be measured using an established assay ^35^,To test this, Δ*galE* cells were grown in LB in the presence or absence of galactose and collected every 30 minutes. Lipids were extracted, labeled with biotin-D-Lys, and detected by immunoblotting. In cells grown in the absence of galactose (lanes with Gal “−”, Fig. 4A-B), accumulation of peptidoglycan precursor was not detected, likely due to their continued use in peptidoglycan synthesis. However, in the presence of galactose, cells accumulated lipid precursors in concentrations easily detected (lanes with Gal “+”, Fig. 4A-B), indicating an inhibition of use of these intermediates. As a positive control, when cells were treated with vancomycin, inhibiting peptidoglycan cross-linking, it resulted in an even stronger accumulation of lipid precursors over untreated cells (Vanc, Fig. 4A-B). Note that in this assay, we did not distinguish whether it is accumulation of lipid I or lipid II.

**Figure 4.**
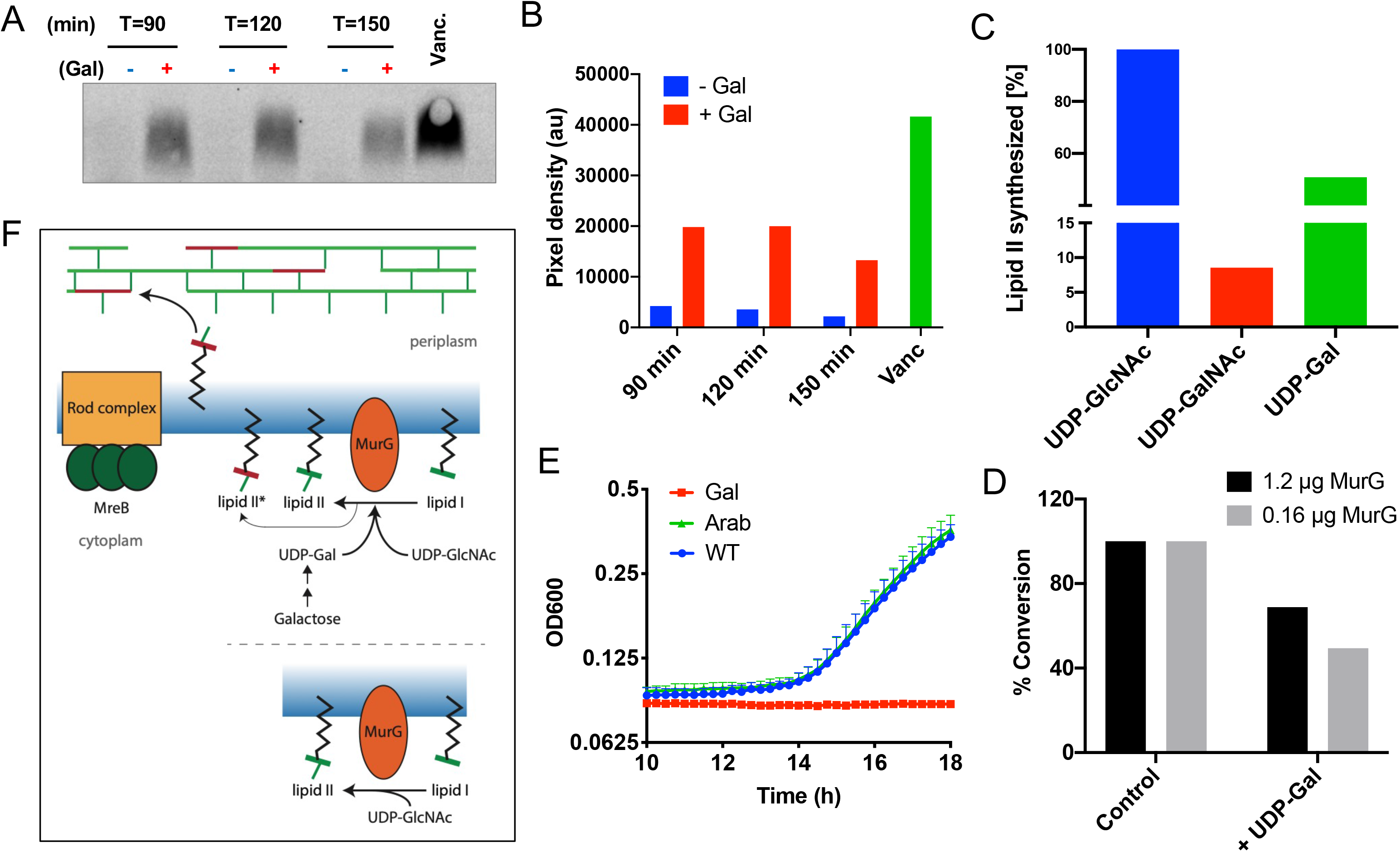
Toxicity is due to block in synthesis of lipid precursors. **(A)** Peptidoglycan lipid precursor levels were measured using a published protocol ^35^. The *galE* mutant (CH62) was grown without (−) and with (+) galactose (0.5%, w/v) in shaking cultures and samples were collected every 30 minutes. Total cell lipids were extracted, biotinylated, and visualized by western blot. As a control, cells treated with cell wall antibiotic vancomycin were similarly assayed. **(B)** The above gel was analyzed, and the amount of lipid I / II quantified by calculating the pixel density using Image J. Assays were repeated three times. A representative result is shown in A and pixel density quantification shown in B. **(C)** To determine if UDP-Gal could be used as a substrate by MurG and incorporated into lipid II, *B. subtilis* MurG was mixed with lipid I and UDP-Gal, UDP-GalNAc, or UDP-GlcNAc, and the amount of resulting lipid II quantified. Upon addition of 5 mM UDP-Gal, there was a 50% production of lipid II as compared to the control of UDP-GlcNAc. Additionally, there was an approximately 7% production of lipid II upon addition of UDP-GalNAc. **(D)** The lipid II yield was measured by using purified *S. aureus* MurG_SA_ and allowing it to react with UDP-GlcNAc without and with the addition of UDP-Gal in an *in vitro* lipid II synthesis assay. Upon addition of UDP-Gal (5 mM) in a ratio of 50:1 to UDP-GlcNAc (100 μM), production of lipid II decreased 50% (at 0.16 μg MurG). The decrease became milder (~30%) if 7.5-fold more purified MurG_SA_ was added (at 1.2 μg MurG). Representative biological replicate shown with error bars representing standard deviation. **(E)** Synergistic lethality of galactose and cell-wall targeting antibiotic vancomycin on WT *S. aureus* cells. The WT *S. aureus* HG003 was grown in 1:4 diluted LB with a sub-MIC concentration (0.2 μg/mL) of vancomycin. Without addition of galactose, growth was slower than normal, due to sub-MIC vancomycin (WT). Upon addition of 0.5% galactose, growth was completely inhibited (Gal). This inhibition was not seen during addition of 0.5% arabinose (Arab). 1:4 diluted LB broth was used here to minimize catabolite repression of galactose metabolism genes in *S. aureus.* **(F)** A cartoon demonstration of how UDP-Gal accumulation inhibits cell wall biosynthesis in bacteria. In the presence of galactose, UDP-Gal is produced and accumulated, which both decreases production of lipid II and produces low levels of toxic lipid II*, which incorporates UDP-Gal in place of UDP-GlcNAc. Lipid II* is unable to be properly polymerized into mature peptidoglycan and causes an “poisoning effect" in peptidoglycan biosynthesis, which leads to eventual cell death.

To further examine the impact of toxicity on MurG *in vivo*, a HALO-MurG fusion protein was created in the Δ*galE* background and engineered cells were imaged by TIRF microscopy. In the absence of galactose, MurG was found to freely diffuse throughout the cell, indicating that it was actively involved in cell wall biosynthesis (-Teixo, Fig. S2A and Supple. Movie 4). The newly discovered antibiotic teixobactin was shown to bind to undecaprenyl-coupled lipid intermediates and thus block cell wall biosynthesis in Gram-positive bacteria ^36^. We first tested if and how this novel antibiotic may impact MurG activities *in vivo*. Upon addition of teixobactin (2.5 μg/mL), free diffusion of HALO-MurG was largely inhibited (+Teixo, Fig. S2A and Supple. Movie 4). As an additional control, the directional movement of MreB filaments was similarly inhibited upon addition of teixobactin (Fig. S2B and Supple. Movie 5). These results suggest that the diffusive state of MurG may be used as an assay for its enzymatic activity. By using this system, we next determined the effect of UDP-Gal accumulation on MurG activities, 0.5% galactose was added to the media and imaging of HALO-MurG performed. By 120 minutes after the addition of galactose, diffusion of HALO-MurG was found to be largely inhibited, further showing that accumulation of UDP-Gal causes an inhibition of MurG activity *in vivo* (Supple. Movie 6).

### UDP-Gal causes production of a “toxic” lipid II *in vitro*

The above evidence suggests that accumulation of UDP-Gal interferes with the function of MurG *in vivo*. To further test this idea *in vitro*, *B. subtilis* MurG was purified from recombinant *E. coli*. MurG protein (1 or 10 μg) was mixed with lipid I and UDP-GlcNAc, UDP-Gal or UDP-GalNAc (at 0.5 or 5 mM). The reaction was allowed to proceed for 60 minutes, quenched, and the amount of lipid II produced quantified by TLC. We found that upon addition of 10 μg of MurG (~8 μM) and 5 mM UDP-Gal, MurG was able to successfully utilize UDP-Gal as a substrate for the generation of lipid II, with a yield approximately 50% that of using the correct ligand UDP-GlcNAc (Fig. 4C and Fig. S3). Importantly, galactose and glucose differ by the orientation of the hydroxyl group at the C4 carbon used in the crosslinking of sugars into mature peptidoglycan (Fig. 1A). Formation of lipid II with UDP-Gal (or UDP-GalNAc) in place of the preferred substrate UDP-GlcNAc would thus produce a “toxic” lipid II and could terminate proper peptidoglycan crosslinking. This result indicates that in addition to inhibition of MurG activity, the toxicity could also be due to formation of “toxic” lipid II generated utilizing UDP-Gal as a substrate, and its “poisoning effect” on peptidoglycan polymerization even though the rate of generating “toxic” lipid II could be much slower than that of the native lipid II.

### UDP-Gal reduces lipid II production by *S. aureus* MurG *in vitro*

We wanted to test if UDP-Gal can similarly interfere with the activity of MurG from another bacterium. To do so, MurG of *Staphylococcus aureus* (MurG_SA_) was expressed and purified from a recombinant *E. coli* strain ^34^. Lipid I was purified using an established biochemical method ^37^ and incubated with purified MurG_SA_ and UDP-GlcNAc. The reaction was allowed to proceed, quenched, and the amount of synthesized lipid II measured. When UDP-Gal was also added in the reaction as an antagonist (in a ratio of 50:1 to UDP-GlcNAc), MurG_SA_ could convert only ~50% of lipid I to lipid II compared to no UDP-Gal addition (0.16 μg MurG_SA_, Fig. 4D). The yield increased to ~70% if more purified MurG_SA_ protein was added (1.2 μg MurG_SA_, Fig. 4D). Intracellular concentrations of UDP-GlcNAc are estimated at approximately 100 μM during the exponential growth ^38^, indicating that the concentration of UDP-Gal would have to reach approximately 5 mM (50 fold in excess) to have an 50% inhibitory effect on MurG, a high concentration, but possible in cells exhibiting toxicity (in which a strong buildup of UDP-Gal is expected). These findings indicate that UDP-Gal can directly interfere the activity of MurG_SA_ at elevated concentrations.

### Galactose increases sensitivity of the wild type *S. aureus* to antibiotics targeting cell wall

Although accumulation of UDP-Gal during galactose metabolism can occur in the *galE* mutant, this nucleotide sugar does not accumulate in, nor does it impact, in any noticeable fashion, the WT cells of *B. subtilis*. Galactose metabolism also does not show any toxicity in *S. aureus* HG003, a model pathogenic strain for clinical infections (data not shown). Interestingly however, when HG003 was grown in the presence of both 0.5% (w/v) galactose and the cell wall targeting antibiotic vancomycin (at a sub-MIC concentration of 0.2 μg/mL), growth was completely inhibited (Gal, Fig. 4E). Inhibition was not observed when galactose was replaced with the same amount of arabinose (Arab, Fig. 4E). In the absence of either sugar, mildly slower than normal growth was recorded due to the presence of sub-MIC vancomycin (WT, Fig. 4E). One possible explanation for the above synthetic lethality is that upon cell wall stress due to sub-MIC vancomycin treatment, the transient accumulation of UDP-Gal from galactose metabolism started to trigger toxicity even in WT cells, due to synergistic inhibitory actions on cell wall biosynthesis.

### *murG* overexpression rescues a similar toxicity phenotype in *Vibrio cholerae*

To see if our findings extend to Gram-negative species as well, a *galE* deletion mutation was constructed in *Vibrio cholerae*. When the *V. cholerae* Δ*galE* cells were grown in LB with galactose (0.5%, w/v), an intermediate toxicity phenotype was observed, judged by a modest decline in cell optical density (red line, Fig. 3E). We were interested in testing if overexpression of the native *murG* gene in *V. cholerae* could give a similar toxicity rescue seen in *B. subtilis*. Indeed, the result shown in Fig. 3E suggests that in *V. cholerae*, overexpression of the native *murG* gene exerted a toxicity rescue phenotype, similar to what we observed in *B. subtilis*. This result indicates that the toxicity-associated bacterial galactosemia could have a common mechanism in distantly related bacteria.

### A suppressor screen on reduced toxicity identifies additional potential targets

To characterize the additional targets or pathways negatively affected in cells exhibiting toxicity in an unbiased fashion, we performed a suppressor screen. A second copy of *galK* and *galT* genes was introduced into the *B. subtilis* Δ*galE* mutant to avoid suppressor mutations in *galK* or *galT* that occurred at a very high frequency in previous studies ^7,39,40^. Cells were grown on LB agar supplemented with 1% (w/v) galactose. Suppressor colonies growing on the plates implied reduced toxicity. They were picked and further verified. A total of 12 suppressor mutants that showed various degrees of toxicity rescue when growing in excess galactose were subjected to whole genome sequencing to characterize the suppressor mutations (Table 1).

**Table 1.**
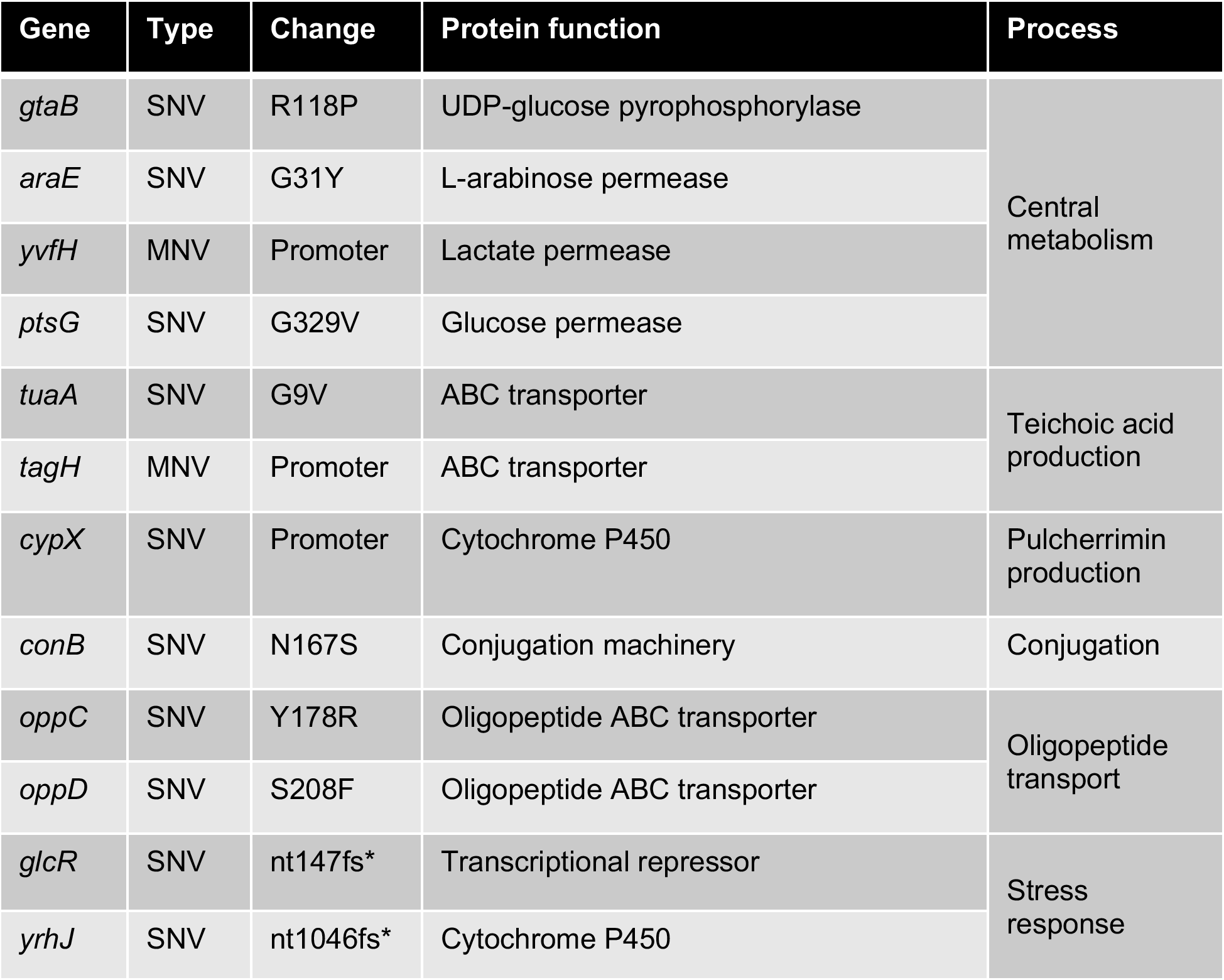
A suppressor screen yields identified mutations in additional putative targets that allowed reduced toxicity. A suppressor screen was performed to identify additional genes or pathways involved in galactose associated toxicity. Δ*galE* cells with a second copy of *galK* and *galT* (YC814) were grown overnight in LB containing 1% (w/v) galactose and allowed to incur natural mutations which reduced toxicity. Mutant strains were enriched, verified, and the suppressor mutations identified by whole genome sequencing. A selection of genes with suppressor mutations, the type of mutations, amino acid sequence change, and gene information are described. fs, frame-shift mutation; SNV, single nucleotide variation; MNV, multiple nucleotide variations.

| Gene | Type | Change | Protein function | Process |
| --- | --- | --- | --- | --- |
| *gtaB* | SNV | R118P | UDP-glucose pyrophosphorylase | Central metabolism |
| *araE* | SNV | G31Y | L-arabinose permease |  |
| *yvfH* | MNV | Promoter | Lactate permease |  |
| *ptsG* | SNV | G329V | Glucose permease |  |
| *tuaA* | SNV | G9V | ABC transporter | Teichoic acid production |
| *tagH* | MNV | Promoter | ABC transporter |  |
| *cypX* | SNV | Promoter | Cytochrome P450 | Pulcherrimin production |
| *conB* | SNV | N167S | Conjugation machinery | Conjugation |
| *oppC* | SNV | Y178R | Oligopeptide ABC transporter | Oligopeptide transport |
| *oppD* | SNV | S208F | Oligopeptide ABC transporter |  |
| *glcR* | SNV | nt147fs* | Transcriptional repressor | Stress response |
| *yrhJ* | SNV | nt1046fs* | Cytochrome P450 |  |

We identified a missense mutation in *gtaB* (R118P, Table 1). The R118 residue is located next to a highly conserved region (^108^KGLGHAVWCA^117^), which maps to a substrate binding helix essential for catalytic activity in the homologous protein of GtaB in *E. coli* (sharing 45% identity in amino acid sequence) ^41^. This may further support our finding of the necessity of GtaB activity for toxic UDP-Gal accumulation (Fig. 2B). Suppressor mutations were also found within genes or in regulatory regions of the genes for known sugar or small molecule transport systems: *araE*, *yvfH*, *ptsG*, *oppC*, *oppD, tagH,* and *tuaA*. Among them, *araE* encodes an arabinose permease. It is proposed that *B. subtilis* does not possess a dedicated galactose permease and galactose uptake relies on the arabinose permease ^42^. Likewise, the hit in *araE* may serve as another validation of the screen. *yvfH* was characterized as a _L_-lactate permease in our previous study ^43^. *ptsG* encodes the PTS system glucose-specific EIICBA component ^44^. *oppC* and *oppD* encode the components of an oligopeptide permease in *B. subtilis* ^45^. A frame shift mutation was also found in the *glcR* gene, which very likely caused loss of function of GlcR. GlcR is a transcription repressor for carbon catabolite control in *B. subtilis*. A deletion mutation in *glcR* caused derepression of certain carbon catabolite control-repressed genes ^46^. Suppressor mutations were also found within a number of cell envelope related genes such as those responsible for wall teichoic acid production (*tagH* and *tuaA*)^47^, as well as those related to cell wall stress response (*cypX* and *yrhJ*)^48^. In many of the above cases, how the suppressor mutations impact the activity of the encoded proteins and how their altered activities allow reduced toxicity is unclear and will require further investigation.

## Discussion

Bacterial galactosemia was discovered five decades ago, but the mechanism causing toxicity was unclear. Here, we provide evidence for the molecular mechanism of toxicity. We found that metabolism of exogenous galactose in the *B. subtilis* Δ*galE* mutant causes inhibition of cell wall biosynthesis, as determined by MreB filament presence and movement. We show that this toxicity is not caused by inhibition of the essential upstream biosynthetic pathway producing UDP-GlcNAc, instead, accumulation of UDP-Gal can directly interfere with the normal function of MurG. This interference includes not only inhibition of MurG, observed *in vivo* by imaging of protein diffusion, but possibly the ability of UDP-Gal to be utilized as an unintended substrate by MurG in the generation of a “toxic” lipid II unable to be polymerized into mature peptidoglycan (Fig. 4F). We find this mechanism may be shared in *V. cholerae* where overexpression similarly produces a partial rescue, as well as in *S. aureus*, where we present evidence that UDP-Gal is able to directly inhibit MurG activity *in vitro*.

Previous work has shown that while the production of lipid II is essential, MurG can bind a number of substrates other than UDP-GlcNAc, a property that has been used as the basis of screen for inhibitory compounds of MurG ^49,50^. This adds to the possibility of nucleotide sugars such as UDP-Gal as inhibitors or as unintended substrates, producing toxic lipid II. The fact that MurG carries out the last cytoplasmic reaction in the peptidoglycan synthesis pathway may add additional lethality if this target is interfered with. To our disappointment, we were unable to identify suppressor mutations in *murG* in our suppressor screen. Likewise, a targeted approach by using site-directed mutagenesis on selected substrate-binding and catalytic residues in *murG* aiming to identify such suppressor mutations was also performed but failed to yield positive results (Habib, unpublished). This may be due to the highly essential nature of MurG, making a mutation which further increases specificity for UDP-GlcNAc while still maintaining enzymatic function, impossible.

Given 1) the near ubiquity of peptidoglycan biosynthesis in bacteria, 2) interference of *S. aureus* MurG-mediated lipid II biosynthesis by UDP-Gal *in vitro*, and 3) the similar toxicity rescue seen in *V. cholerae*, it is likely that this mechanism of toxicity is conserved in various bacterial species. It is also possible that this inhibition is not unique to MurG, but to other essential proteins involved in the utilization of UDP-GlcNAc that may have active site chemistry similar to MurG ^49^. Galactosemia was discovered in human century ago ^51^. Apparently human cells do not have peptidoglycan and MurG. However, they possess glycocalyx on the cell surface, which functions in cell–cell recognition, communication and adhesion, as well as other functionally important glycoproteins. The synthesis of glycocalyx depends on glycosyltransferases that are functional analogs to bacterial glycosyltransferases such as MurG ^52^. It will be interesting to know if the activity of any of those essential glycosyltransferases is altered in the galactosemia patients, though investigation like this is outside the scope of this study.

Finally, in WT *B. subtilis*, galactose metabolism does not cause toxicity. It is conceivable that when WT cells metabolize galactose, accumulation of UDP-Gal seldomly reaches the threshold that triggers toxicity. As discussed earlier, UDP-Gal production is necessary for several important cellular processes involving various polysaccharide syntheses. Thus, there must be a homeostatic regulation for UPD-Gal production in the bacterium. Many of these nucleotide sugars and phosphorylated monosaccharides, if accumulated above certain levels, could create an adverse impact on cell physiology or toxicity. Understanding the homeostatic regulation of those sugar intermediates and impact under mis-regulation may reveal new cellular targets and provide new ideas of antibacterial solutions. The *galE* mutant used in this study can be considered an unusual case, in which there is misregulation of specific sugar metabolites. Under this misregulation, we discovered that the cytoplasmic stages of cell wall biosynthesis can indeed be the targets of sugars native to the cells. Using such an artificial system conveniently allows us to understand the key points of regulation as well as potential novel antibacterial ideas. Finally, although galactose metabolism does not cause any toxicity in WT cells, the increase in sensitivity to cell-wall targeting antibiotics exhibited in the WT *S. aureus* upon simultaneous galactose metabolism demonstrates possible clinical relevance of galactose metabolism-induced cellular stress or toxicity.

## Materials and Methods

### Strains, medias, and reagents

*B. subtilis* and *E. coli* strains were routinely cultured in lysogeny broth (LB). All strains used in this study are listed in Table S1. Insertional deletion strains, if not specified in the methods, were purchased from the Bacillus Genome Stock Center (BGSC, Columbus, OH). Insertional deletion mutations were then introduced into the *B. subtilis* NCIB3610 background by transformation. MSgg media for biofilm formation was prepared according to the published protocol ^53^. To induce the toxicity, cells were grown in LB (1% tryptone, 0.5% yeast extract, and 1% NaCl) with the addition of 0.5% (w/v) galactose and 0.01% (w/v) arabinose. Addition of arabinose is to induce the *araE*-encoded arabinose permease for galactose uptake ^42^. If required, antibiotics were added at the following concentrations for *B. subtilis*: 10 μg ml^−1^ kanamycin, 50 μg ml^−1^ spectinomycin, and 5 μg ml^−1^ tetracycline. For *E. coli* strains, when required, antibiotics were used at: 100 μg ml^−1^ ampicillin and at 50 μg ml^−1^ kanamycin. Chemicals, including galactose (CAS No. 59-23-4), arabinose (CAS No. 5328-37-0), UDP-NAc-Glucosamine (CAS No. 91183-98-1), and UDP-Galactose (CAS No. 137868-52-1) were purchased from Sigma-Aldrich. Restriction enzymes were purchased from New England Bio-labs (NEB, MA, USA). Oligonucleotides (sequences provided in Table S2) were purchased from Integrated DNA Technologies (IDT, IA, USA). DNA sequencing was performed at Genewiz (NJ, USA).

**Strain construction** was described in Supplementary Materials.

### Biofilm development

For pellicle biofilm development, the Δ*gtaB* cells were inoculated from colonies on an overnight LB agar plate into 3 mL of LB broth and grown with shaking at 37°C to log phase. Cells were then subcultured 1:1000 into 7 mL of MSgg in a 6-well polyvinyl plate (VWR, PA, USA). Plates were incubated in static conditions at 30°C for 3 days and were then imaged using a Leica MZ10F macroscope and Leica DMC2900 camera. For colony biofilm development, Δ*gtaB* cells were similarly prepared and 3 ul of the log phase cells was spotted onto the MSgg media solidified with 1.5% agar (w/v). Plates were incubated at 30°C for 3 days and were then imaged using a Nikon Coolpix camera.

### Phase contrast microscopy

Phase contrast and fluorescent imaging for live/dead staining was performed on a Leica DM5000B light microscope. Cells were grown to mid-exponential phase and collected by centrifugation for 1 minute at 16,000 g, resuspended in phosphate buffered solution (PBS), and concentrated 10-fold. 2 μL of the resuspended culture was spotted on a glass slide and covered with poly-L-lysine coated coverslip. Resulting slides were viewed at 100x magnification and images were captured with Leica DFC300G camera. To perform live / dead staining, cells were collected, concentrated as above, and treated with live/dead staining dyes (Invitrogen, Carlsbad, CA) according to manufacturer’s protocol with resulting samples imaged under GFP (for live cells) and Texas Red (for dead cells) fluorescent filters. The ratio of live/dead cells was manually counted for about a total of 300 cells.

### Scanning Electron Microscopy

*B. subtilis* Δ*galE* cells were grown overnight in LB to stationary phase and subcultered 1:100 in LB supplemented with galactose and were grown 3 hours shaking at 37°C. Cells were collected via centrifugation at 5,000 g and washed twice with PBS. Cells were allowed to adhere to a glass coverslip treated with poly-L-lysine for 1 hour. Adhered cells were fixed in a solution containing 2.5% glutaraldehyde and 0.1M sodium cacodylate at pH 7.2 for 1 hour and then rinsed for 10 minutes in 0.1 sodium cacodylate buffer pH 7.2 three times. Cells were infiltrated with 1% osmium tetroxide in 0.1M sodium cacodylate pH 7.2 for 1 hour and then washed again three times with 0.1 sodium cacodylate buffer pH 7.2 for 10 minutes each wash. Cells were then dehydrated in ethanol at 30%, 70%, 85%, and 95%, for 10 minutes, and finally in 100% ethanol for three times at 10 minutes each wash. Samples were then dried in a critical point dryer utilizing carbon dioxide, mounted to specimen mounts, using double-sided carbon adhesive tape, and sputter-coated with 5nm platinum. Samples were then examined using a high-resolution field emission SEM Hitachi S-4800 under vacuum using an acceleration voltage of >2.0 kV.

### Single-cell growth rate time lapse experiments

2-μL culture was spotted on No. 1.5 glass-bottomed dishes (MatTek, MA) under 3% agarose pads made with casein hydrolysate (CH) medium. The phase-contrast images were collected on a Nikon Ti microscope equipped with a Hamamatsu ORCA Flash4 CMOS camera with a Nikon Plan Apo λ 100×/1.4NA objective. Images were acquired from ~50 fields of view every 2 min for a total of 2-3 hour using NI Element. The phase-contrast time-lapse movies were analyzed using a custom-built package in MATLAB.

### Imaging of MreB-mNeonGreen by TIRF microscopy

Cells were placed under a CH agarose pad with galactose (0.5%, w/v) on a glass coverslip for imaging. Images were collected on a Nikon Ti microscope equipped with a Hamamatsu ORCA Flash4 CMOS camera with a Nikon 100X NA 1.45 objective. Exposure time is 300ms and the time interval 1 second. A 488nm laser line is used for the imaging of mNeonGreen. The method used to analyze the density of MreB filaments has been published previously ^28^. First the phase images were segmented, and the fluorescence time-lapses were analyzed based on the segmentation mask of the phase image. Next for each cell the kymographs were generated for each row of pixels along the midline of the cell, and the time-lapse movie was converted into a single 2D image. Then the MreB filament were registered in the 2D image. The MreB filament density is normalized to the density in WT cells (YS09). All of the image analyses were performed using custom MATLAB code.

### Imaging of HaloTag-MurG by TIRF microscopy

HaloTag-MurG cells were incubated with 50 pM of HaloTag-JF549 ligand for 30 minutes and then washed twice to get rid of free dyes. Images were collected on a Nikon Ti microscope equipped with a Hamamatsu ORCA Flash4 CMOS camera with a Nikon 100X NA 1.45 objective. Exposure time is 300ms and the time interval 1 second. A 561nm laser line is used for the imaging of JF549-HALO dye.

### Bulk growth analysis

For bulk culture growth measurements by cell optical density (OD_600_), cells were grown overnight in LB plus applicable antibiotics to stationary phase with shaking at 200 rpm at 37°C. Cells were subcultured 1:100 in respective media and grown to mid exponential phase (OD_600_ = 0.4-0.5). Cells were subcultured 1:100 in 96-well tissue culture plates with 170 μL final volume of media in each well. Plates were read by BioTek plate reader with shaking at 200 rpm and incubating at 37°C with OD_600_ readings taken even 15 minutes. Experiments were performed at least three times and representative data set were shown.

To determine the effect of galactose metabolism on growth of *Staphylococcus aureus*, cells were grown shaking at 200 rpm at 37°C overnight. Cells were subcultured 1:200 in LB diluted 1:4 with sterile water. Vancomycin was added to all samples at 0.2 μg/mL, and galactose or arabinose was added at 0.5% (w/v), with sterile water used as a control. Samples were grown in a 24-well tissue culture plate with 2mL final volume in each well and read as above.

### Peptidoglycan precursor measurement

Measurement of peptidoglycan precursor was performed using published methods with minor modifications ^35^. Briefly, *B. subtilis* strain CH62 was grown overnight in LB with tetracycline to stationary phase and subculture 1:100 into both LB and LB plus 0.5% (w/v) galactose. Vancomycin was used as a control at 10 μg mL^−1^ added at OD_600_ =0.4 and were treated for 30 minutes before collection. 10 mL of cells from both conditions were collected every 30 minutes and spun down for 10 minutes at 16,000 g. Total lipid content of the cells was extracted via two-phase system and concentrated *in vacuo.* The resulting extract was mixed with biotinylated lysine (BDL) probe and purified PBP4 protein and allowed to react for an hour at 25°C before being quenched. Reaction mixture was loaded on to a 15% SDS-PAGE and transferred to Immuno-Blot PVDF membrane. BDL was detected utilizing HRP-Conjugated Streptavidin antibody and imaged on BioRad ChemiDoc MP. The resulting gel was quantified for pixel density using ImageJ. Experiment was performed three times and a representative image is presented.

### MurG *in vitro* assay

MurG activity using different UDP-activated substrates was tested as previously described with minor modifications ^34^. Briefly, MurG activity assays were performed in a final volume of 30 μL containing 3 nmol of purified lipid I, 5.8 to 50 mM of either UDP-Gal or UDP-GalNAc in 60 mM Tris-HCl, 5 mM MgCl2, pH 7.5, and 0.5% Triton X-100 in the presence of 1 or 10 μg (~0.8 or 8 μM) of purified MurG_Bsub_-His_6_ proteins. The reaction mixture was incubated for 60 min at 30 °C. Lipid intermediates were extracted with an equal volume of n-butanol/pyridine acetate, pH 4.2 (2:1; v/v) and analyzed by thin layer chromatography (TLC) using chloroform/methanol/water/ammonia (88:48:10:1, v/v/v/v) as the solvent ^34^. The quantitative analysis of lipid II extracted to the butanol phase was carried out using ImageJ ^54^. The inhibitory effect of UDP-Gal (molar ratio of 50:1 with respect to UDP-GlcNAc) on MurG_SA_ activity was tested in a final volume of 30 μL containing 1 nmol of purified lipid I, 10 nmol of UDP-GlcNAc, 0.1 nmol [14C]UDP-GlcNAc (1.23 kBq) in 60 mM Tris-HCl, 5 mM MgCl2, pH 7.5, and 0.5% Triton X-100 in the presence of 0.16 or 1.2 μg of purified MurG-His_6_ protein. The reaction mixture was incubated for 30 min at 30 °C. Lipid intermediates were extracted and analyzed by thin layer chromatography (TLC) as described above. Radiolabeled lipids were visualized by phosphorimaging using a Storm imaging system and quantification was performed using the software ImageQuant TL (GE Healthcare),

**Whole genome sequencing** was described in Supplementary Materials.

## Acknowledgments

We would like to thank Luke Lavis for JF dyes, and Dr. Kim Lewis (Northeastern University) for the gift of teixobactin. This work was supported by a grant from the National Science Foundation (MCB1651732) to YC. This work was also funded by National Institutes of Health Grants DP2AI117923-01 to EG, as well as support from the Volkswagen Foundation. Funding was also provided by the German Research Foundation (Project-ID 398967434 – TRR261 to T.S. and A.M) and an NIH grant (AI120489) to JZ.

## Author contributions

C.H., Y.S., E.G., and Y.C. conceived the project; C.H. and Y.S. cloned the strains; C.H. A.M., T.S., M.L., and Y.S. planned and performed the experiments; C.H., A.M., Y.S., and Y.C. analyzed the data; Y.S. wrote the code for the single cell analyses; C.H., Y.S., and Y.C. wrote the manuscript, with contributions and editing from all authors.

## Supplemental files_Habib et al.

## Supplemental Methods

### Strain construction

General methods for molecular cloning followed published protocols ^1^. Long-flanking PCR mutagenesis was used to generate insertional deletion mutations ^2^. To construct the *B. subtilis* Δ*galE* strain, genomic DNA was isolated from the strain YC814 and transformed into WT NCIB 3610 (hereafter 3610) and plated on LB agar plates under tetracycline selection ^3^. To construct the *gtaB* deletion strain (YC876), the insertional deletion mutation in *gtaB* was created by long-flanking PCR mutagenesis. The four primers (delta-gtaB-P1 to delta-gtaB-P4) used for *gtaB* PCR mutagenesis are listed in Table S2. The double mutant of Δ*galE*Δ*gtaB* was made by introducing Δ*galE*::*tet* from YC814 to YC876 by transformation and selecting the transformants on LB plus both tetracycline and kanamycin. To construct an IPTG inducible copy of *gtaB,* the native *gtaB* gene was amplified by PCR using primers P_gtaB_-F1 and P_gtaB_-R1 and isolated 3610 genomic DNA as the template. The purified PCR product was cloned into HindIII and SalI restriction sites of the plasmid pDR111. The resulting plasmid was transformed into and purified from *E. coli* strain DH5⍺, verified by DNA sequencing, and then transformed into the *B. subtilis* lab strain PY79. Transformants were selected for double-crossover recombination at the chromosomal *amyE* locus on LB agar plate with spectinomycin. The genomic DNA of successful transformants was purified and used to transform the Δ*galE*Δ*gtaB* double mutant of *B. subtilis* outlined above. The generation of IPTG inducible *murG* was performed as above, amplified using primers P_murG_-F1 and P_murG_-R2 and utilizing the HindIII and NheI restriction sites of pDR111. Similarly, the IPTG inducible copy of *tagB*, *murAA*, and *mnaA* were performed utilizing respective primers tagB-F1 and tagB-R1, murAA-F1 and murAA-R1, and mnaA-F1 and mnaA-R1. To generate the Δ*galE/mreB*-*mNeonGreen* reporter strain, genomic DNA from YC814 was isolated and transformed into bYS09, containing a native promoter protein fusion of MreB-mNeonGreen ^4,5^, and plated on LB agar plate selecting for erythromycin resistance. To construct the Δ*galE/murG/mreB*-*mNeonGreen* strain, the pDR111-*murG* plasmid constructed above was transformed into the Δ*galE/mreB*-*mNeonGreen* strain and selected for by plating on LB agar with spectinomycin. To generate IPTG inducible copies of *glmU*, *glmM*, *glmS*, and *glmR*, genomic DNA from strains HB16910 (*glmM*), HB16951 (*glmR*), HB21922 (*glmU*), and HB21942 (*glmS*), [gifts of the J. Helmann] were prepared and transformed into a Δ*galE* background selecting for chloramphenicol resistance.

The HALO-murG fusion (bYS993) was generated by transforming PY79 with a Gibson assembly consisting of five fragments: 1) PCR with primers oMD191 and oMD108 and PY79 template genomic DNA; 2) PCR with primers oJM28 and oJM29 and pWX467 [gift of D. Rudner] template genomic DNA (containing the kanamycin-resistance cassette *loxP-kan-loxP*); 3) PCR with primers oMD234 and oMD232 and template pDR150 [gift of D. Rudner] genomic DNA; 4) PCR with primers oYS600 and oAB60 and HALO-tag template; 5) PCR with primers oYS835 and oMD197 and template PY79 genomic DNA. *mreB-mNeonGreen* fusion in a *galE* deletion background (bYS992) was generated by transforming bYS09 with genomic DNA from GalE KO::erm. bYS995 [*amyE::Phyperspank*-*HALO*-*murG::kan*, *galE*::erm] was generated by transforming bYS993 with genomic DNA from *galE::erm*.

*The amyE::Phyperspank-mraY::erm* (bYS424) was generated by transforming PY79 with a Gibson assembly consisting of five fragments: 1) PCR with primers oMD191 and oMD108 and PY79 template genomic DNA; 2) PCR with primers oJM28 and oJM29 and pWX467 [gift of D. Rudner] template genomic DNA (containing the erythromycin -resistance cassette *loxP-erm-loxP*); 3) PCR with primers oMD234 and oMD232 and template pDR150[gift of D. Rudner] genomic DNA; 4) PCR with primers oYS682 and oYS357 and PY79 template; 5) PCR with primers oMD196 and oMD197 and template PY79 genomic DNA.

bYS997 [*mreB-mNeonGreen*, *amyE::Phyperspank-murG::spec*, *galE::erm*] was generated by transforming bYS992 with genomic DNA from *amyE::Phyperspank-murG::spec*. bYS998 [*mreB-mNeonGreen*, *amyE::Phyperspank-MurAA::spec*, *galE::erm*] was generated by transforming bYS992 with genomic DNA from CH125 (bCH125). bYS999 [*mreB-mNeonGreen*, *amyE::Phyperspank-MraY::erm*, *galE::erm*] was generated by transforming bYS992 with genomic DNA from *amyE::Phyperspank-mraY::erm* (bYS424).

Overexpression of the native *E. coli murG* was performed by isolating genomic DNA from the *E. coli* lab strain DH5⍺ and amplifying native *murG* gene utilizing primers P_murG_-F2 and P_murG_-R2. The resulting purified PCR product was cloned into pUC18 plasmid utilizing restriction sites BamHI and PstI. The recombinant plasmid was transformed into and purified from DH5⍺ and sequence verified. The resulting plasmid was transformed into the *E. coli* strain JW0742 (Δ*galE*, the Keio collection)^6^, and selected for by plating on LB agar plate with ampicillin*. B. subtilis* 168 *murG* was amplified using primers murGBsub FW and RV and cloned into pET21b vector (Novagen) by InFusion cloning to generate a C-terminal His6 fusion protein. The recombinant plasmid was transformed into DH5⍺, verified by DNA sequencing, and then transformed into BL21/DE3 for protein expression. Expression of MurGBsub-His6 and MurGSaur-His6 and purification were performed as described ^7^.

*V. cholerae* E1 Tor C6706 ^8^ was used as the wild-type strain in this study. In-frame deletion of *galE* (VCA0774) was constructed by cloning the regions flanking target genes into the suicide vector pWM91 containing a *sacB* counter-selectable marker ^9^ utilizing primers galE-Up-XhoI-5, galE-Up-3, galE-Down-5, and galE-Down-NotI-3. Double-crossover recombination mutants were selected using 10% sucrose plates and confirmed via PCR. Complementation of *murG* was constructed by cloning the complete *murG* (VC2401) sequence into the pSRKTc vector utilizing primers murG-NdeI-5’ and murG-XbaI-3’ ^10^. For *in vitro* growth experiments, cells from overnight cultures grown in LB medium of the WT, the *galE* mutant and *murG* complemented mutant were inoculated 1:100 into fresh LB medium supplemented with tetracycline of 2 μg mL^−1^ and IPTG (isopropyl-beta-D-thiogalactopyranoside) of 0.5 mM at a final concentration, when indicated, 0.05% galactose was included, statically grown at 37℃ and measured OD_600_ using Bioscreen C (Type FP-1100-C) instrument at time points indicated.

### Whole genome sequencing

To prepare DNA for whole genome sequencing, all strains were grown to a mid-exponential phase OD_600_ of 0.4 and 1 mL of culture spun down 2 minutes at 16,000 g. Cells were resuspended in 450 μL of ddH_2_O with 50 μL 500 mM EDTA and 60 μL lysozyme at 20 mg/mL, and incubated for 30 minutes at 37 °C. After incubation, 650 μL of cell lysis solution (Promega) was added and tube was inverted several times, followed by 250 μL of protein perception solution (Promega). Tube was vortexed until homogenous and centrifuged for 5 min at 16,000 g. 900 μL of supernatant was transferred to a new tube and 600 μL isopropanol added to precipitate DNA, with the tube inverted several times and incubated at room temperature for 2 min, then spun 16,000 g for 5 min. DNA pellet was washed in 70% ethanol and spun down at 16,000 g. Tube was dried at room temperature under air flow for 10 minutes and then resuspended in 100 μL of ddH_2_O. 2 μg of DNA was digested for 30 min with dsDNA Fragmentase (NEB) and then stopped with EDTA. DNA fragments of approximately 200-bp were selected for using AMPure XP beads (Beckman Coulter, Indianapolis, IN) according to manufacturer protocols. Resulting fragments were prepared and labeled for whole genome sequencing using NEBNExt Ultra DNA Library Prep Kit (NEB) from Illumina technologies. Determination of resulting DNA library purity and concentration, and subsequent sequencing was performed by the Harvard Bauer Core Facility (Harvard, Cambridge, MA).

## Supplemental Figure legends

**Fig. S1.**
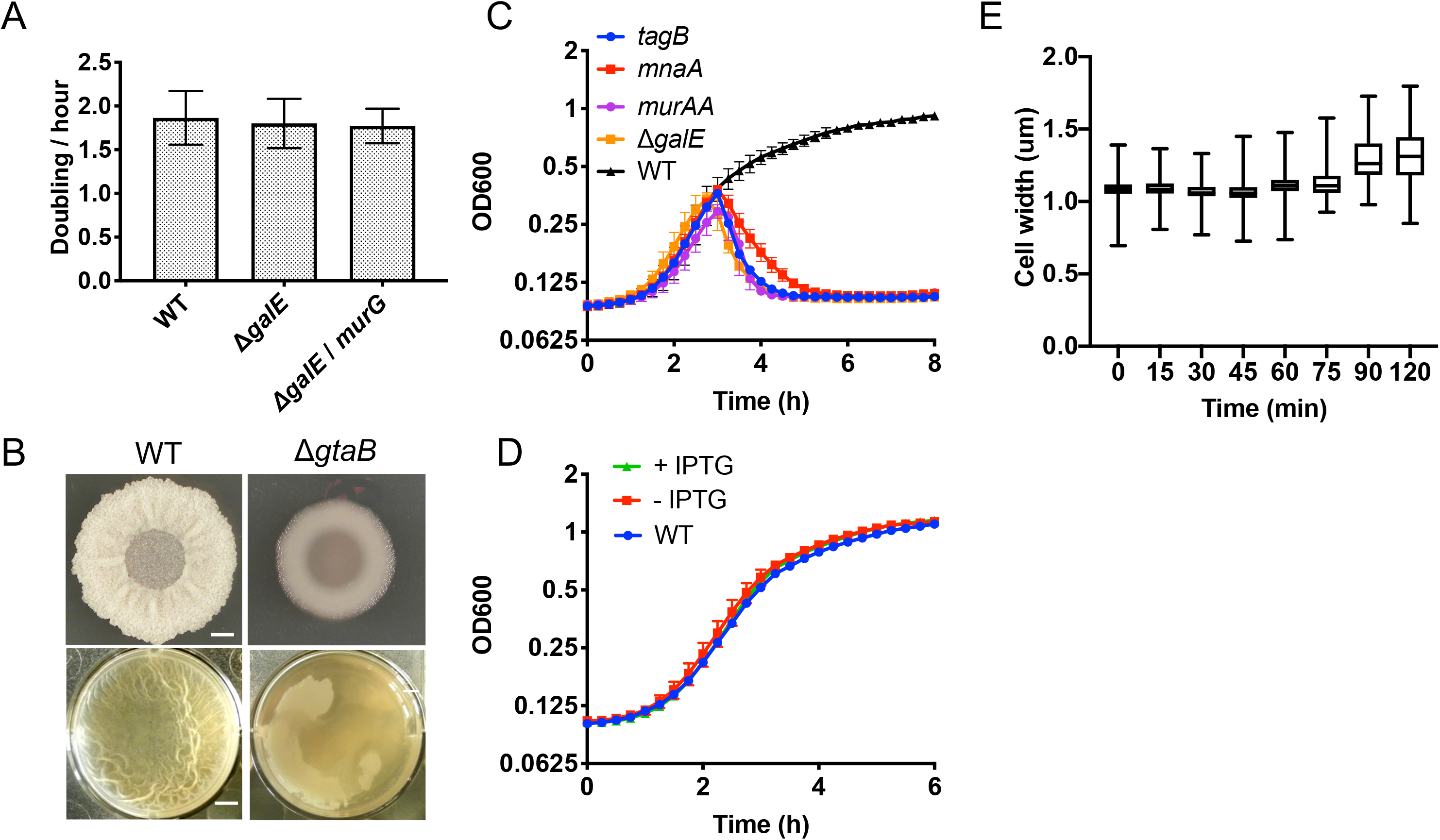
**(A)** To determine if any difference in growth rate was apparent at a single cell resolution upon deletion of *galE*, the same strains were grown on a CH media agar pad and imaged over time. Similar to OD600 measurements, no change in rate was observed. Bars indicate standard deviation. Colony and pellicle biofilm development by the WT (3610) and the Δ*gtaB* mutant (YC876) in solid and liquid MSgg media. The Δ*gtaB* mutant was unable to form robust biofilms, presumably due to lack of the essential nucleotide sugar UDP-Glc for making exopolysaccharides of the biofilm matrix. Biofilms were incubated at 30°C for 3 days before imaging. Scale bars indicate 1 mm, pellicle well diameter is 3.5 cm. **(C)** To determine if the rescue exhibited by *murG* was unique to *murG* or shared among other glycotransferases which utilize UDP-NAG, an IPTG inducible copy of *murAA* (CH125)*, glmS* (CH127)*, tagB* (CH123)*, and mnaA* (CH129) were each respectively introduced into the *B. subtilis* Δ*galE* mutant and (CH062) and grown in LBga shaking at 37°C with 0.5 mM IPTG, along with wild type and Δ*galE* strains, and OD600 measurements taken every 15 minutes. Overexpression of genes had no rescue associated with it as compared to Δ*galE* strain. **(D)** To determine if the rescue seen by *murG* overexpression was due to an increase in growth rate, the Δ*galE/*IPTG*-murG* strain created above as well as wild type was grown shaking at 37°C in LB without (−) and with (+) 500 μM of IPTG with OD600 measurements taken every 15 minutes. Overexpression of *murG* had no effect on growth rate. **(E)** The effect of MurG overexpression on toxicity as a function of maximal cell width of ~100 cells using microscopy. Cells were grown in galactose and MurG overexpression induced using 500μM IPTG. Compared without overexpression of MurG, cells were able to maintain a consistent max cell width for a greater amount of time, indicating a rescue in toxicity resulting from MurG. Error bars represent standard deviations.

**Fig. S2.**
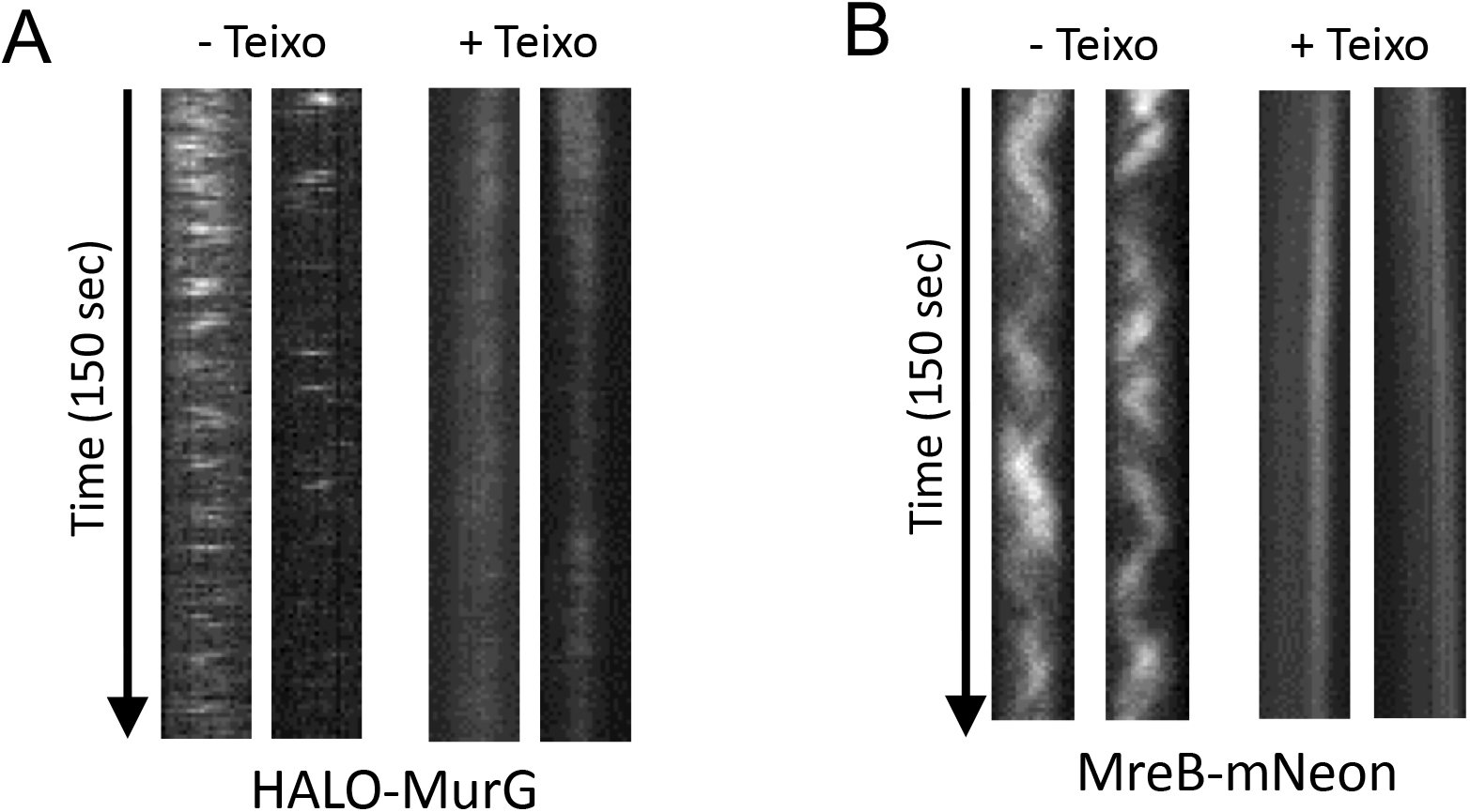
**(A)** *B. subtilis ΔgalE* mutant expressing a HALO-MurG fusion was imaged using TIRF microscopy on a CH agar pad with or without the addition of 25 ug/mL of teixobactin. Kymographs were created utilizing the images obtained. It was found that upon addition of teixcobactin, activity of MurG was eliminated, suggesting that the diffusion of MurG could serve as a reporter for its activity. As a control, the *B. subtilis ΔgalE* mutant expressing the MreB-mNeonGreen fusion was similarly imaged with and without the addition of 25 ug/mL of teixobactin. Upon addition, MreB diffusion was also arrested, providing additional support for the use of movement as a measurement for protein activity.

**Fig. S3.**
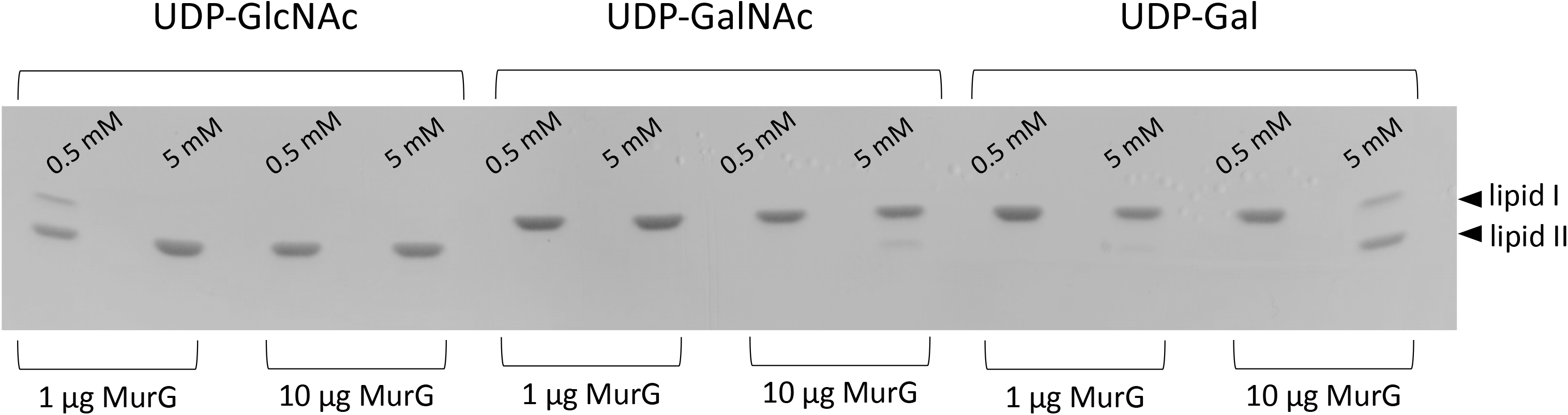
One or ten microgram (ug) of B. subtilis MurG was mixed with either 0.5 or 5 mM of UDP-GlcNAc, UDP-GalNAc, or UDP-Gal (respectively) to determine the ability of MurG to utilize these substrates in the generation of lipid II from lipid I. The products of the reactions were run on TLC and quantified, showing that MurG can utilize both UDP-GalNAc as well as UDP-Gal as substrates, though at a reduced efficiency to the preferred substrate of UDP-GlcNAc

**Supplemental movie 1.** The *B. subtilis ΔgalE* mutant expressing a MreB-mNeonGreen fusion protein in native locus was imaged via TIRF microscopy after the addition of galactose (0.5%, w/v) onto a CH agar pad. By 60 minutes after galactose addition, most of MreB filaments completely ceased movement or were depolymerized, indicating inhibition of cell wall biosynthesis in cells exhibiting strong toxicity. Cells were imaged at 1 second intervals with an exposure time of 300ms.

**Supplemental movie 2.** The *B. subtilis ΔgalE* mutant expressing a MreB-mNeonGreen fusion protein in native locus and Phyperspank MurG in AmyE locus was imaged via TIRF microscopy after the addition of galactose (0.5%, w/v) and 1mM IPTG onto a CH agar pad. By 90 minutes after galactose addition, directional motion of MreB filaments could still be observed. Cells were imaged at 1 second intervals with an exposure time of 300ms.

**Supplemental movie 3.** The *B. subtilis ΔgalE* mutant expressing a MreB-mNeonGreen fusion protein in native locus and Phyperspank MurAA, MraY or MurG in AmyE locus was imaged via TIRF microscopy after the addition of galactose (0.5%, w/v) and 1mM IPTG onto a CH agar pad. After 90 minutes galactose addition, directional motion of MreB filaments could only be observed for cells with MurG overexpression but not for cells with MurAA or MraY overexpression. Cells were imaged at 1 second intervals using TIRF microscopy with an exposure time of 300ms.

**Supplemental movie 4.** *B. subtilis ΔgalE* mutant expressing a HALO-MurG fusion was imaged using TIRF microscopy on a CH agar pad without or with the addition of teixobactin (2.5 and 25 ug/mL). Cells were imaged at 1 second intervals using TIRF microscopy with an exposure time of 500ms.

**Supplemental movie 5.** *B. subtilis* expressing a MreB-mNeonGreen fusion was imaged using TIRF microscopy on a CH agar pad without or with the addition of teixobactin (2.5 and 25 ug/mL). Cells were imaged at 1 second intervals with an exposure time of 300ms.

**Supplemental movie 6.** To determine the effect of UDP-Gal accumulation on MurG *in vivo*, the *B. subtilis ΔgalE* mutant expressing a HALO-MurG fusion was imaged via TIRF microscopy 60, 90, and 120 minutes after the addition of galactose (0.5%, w/v) onto a CH agar pad. After 90 minutes after galactose addition, a significant population of MurG had arrested movement, indicating that UDP-Gal accumulation causes an inhibition of MurG activity. Cells were imaged at 1 second intervals with an exposure time of 500ms.

**Table S1:** Strains used in this study.

| Strains | Genotypes | References |
| --- | --- | --- |
| *B. subtilis* PY79 | A laboratory strain for genetic manipulation | <sup>11</sup> |
| *B. subtilis* NICB3610 | Undomesticated *B. subtilis* strain capable of biofilm formation | <sup>12</sup> |
| CH062 | $\Delta galE::tet^R$ in 3610 | this study |
| CH112 | *amyE::P<sub>hyspank</sub>-murG*, $\Delta galE::tet^R$ in 3610 | this study |
| CH117 | $\Delta gtaB::spec^R$, $\Delta galE::tet^R$, *amyE::P<sub>hyperspank</sub>-gtaB* in 3610 | this study |
| CH123 | $\Delta galE::tet^R$, *amyE::P<sub>hyspank</sub>-tagB::spec^R* in 3610 | this study |
| CH125 | $\Delta galE::tet^R$, *amyE::P<sub>hyspank</sub>-murAA::spec^R* in 3610 | this study |
| CH127 | $\Delta galE::tet^R$, *amyE::P<sub>hyspank</sub>-glmS::spec^R* in 3610 | this study |
| CH129 | $\Delta galE::tet^R$, *amyE::P<sub>hyspank</sub>-mnaA::spec^R* in 3610 | this study |
| CH143 | $\Delta galE::kan^R$, pUC18-*murG* in DH5a | this study |
| CH150 | $\Delta galE::tet^R$, *amyE::P<sub>hyspank</sub>-glmU* | this study |
| CH151 | $\Delta galE::tet^R$, *amyE::P<sub>hyspank</sub>-glmM* | this study |
| CH152 | $\Delta galE::tet^R$, *amyE::P<sub>hyspank</sub>-glmS* | this study |
| CH153 | $\Delta galE::tet^R$, *amyE::P<sub>hyspank</sub>-glmR* | this study |
| YC814 | $\Delta galE::tet^R$, *amyE::P<sub>hyspank</sub>-galT-galK::spec^R* in 3610 | <sup>13</sup> |
| YC876 | $\Delta gtaB::kan^R$ in 3610 | this study |
| DH5α | A laboratory strain of *E. coli* | Invitrogen |
| VCA0774 | $\Delta galE::tet^R$ in *V. cholerae* E1 Tor C6706 | this study |
| VC2401 | $\Delta galE::tet^R$, pSRKTc-*murG* in *V. cholerae* E1 Tor C6706 | this study |
| bYS09 | *mreB-mNeonGreen* | <sup>5</sup> |
| bYS424 | *amyE::Phyperspank-mraY::erm* | this study |
| bYS992 | *mreB-mNeonGreen*, $\Delta galE::erm$ | this study |
| bYS997 | *mreB-mNeonGreen*, $\Delta galE::erm$, *amyE::P<sub>hyperspank</sub>-murG::spec* | this study |
| bYS998 | *mreB-mNeonGreen*, $\Delta galE::erm$, *amyE::P<sub>hyperspank</sub>-murAA::spec* | this study |
| bYS999 | *mreB-mNeonGreen*, $\Delta galE::erm$, *amyE::P<sub>hyperspank</sub>-mraY::erm* | this study |
| bYS993 | *amyE::P<sub>hyperspank</sub>-HALO-murG::kan* | this study |
| bYS995 | *amyE::P<sub>hyperspank</sub>-HALO-murG::kan*, $\Delta galE::erm$ | this study |

**Table S2:**
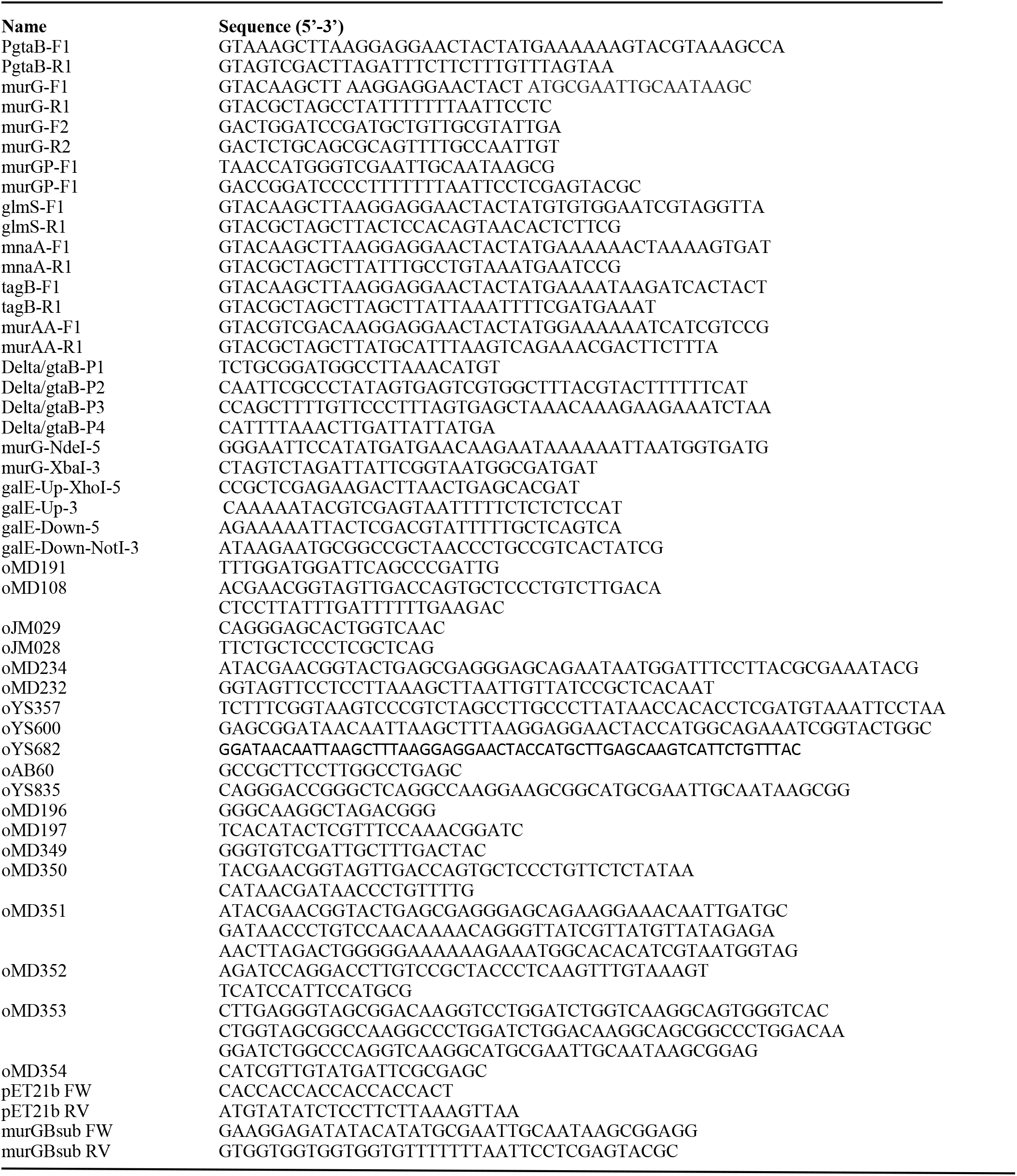
Oligonucleotides used in this study.

